# Passive Acoustic Monitoring within the Northwest Forest Plan Area: 2025 Annual Report

**DOI:** 10.64898/2026.05.26.727992

**Authors:** Damon B. Lesmeister, Julianna M. A. Jenkins, Zachary J. Ruff, Natalie M. Rugg, Simeon Abidari, Sallie Beavers, Roger Christophersen, Raymond J. Davis, Miranda Terwilliger, Aaron Henderson, Brandon Henson, Julia Kasper, Christopher McCafferty, Dave Press, Suzanne Reffler, James K. Swingle, Alaina D. Thomas, Kirsten Wert

## Abstract

The Northwest Forest Plan (NWFP) Passive Acoustic Monitoring (PAM) program is a large-scale interagency biodiversity monitoring framework designed to assess the status and trends of northern spotted owls (*Strix occidentalis caurina*), barred owls (*Strix varia*), marbled murrelets (*Brachyramphus marmoratus*), and broader forest biodiversity across federally administered lands in the Pacific Northwest. In 2025, we deployed autonomous recording units at 2,095 sampling stations within 532 5-km^2^ hexagons randomly selected across approximately 24 million acres of federal forest lands in Washington, Oregon, and California. These deployments generated 1.37 million hours of acoustic recordings that we processed using the convolutional neural network PNW-Cnet v5 for automated species identification and subsequent human validation of focal species detections. Northern spotted owls were detected in 28% of sampled hexagons range-wide, including 16% in Washington, 25% in Oregon, and 60% in California. Occupancy patterns remained consistent compared to previous years, with higher occupancy concentrated in southern portions of the geographic range and continued low occupancy across much of the Washington Cascades and Oregon Coast Range. Barred owls were detected in 86% of sampled hexagons and remained broadly distributed throughout most of the NWFP area. Marbled murrelets were detected in 50% of reviewed hexagons within NWFP marbled murrelet management zones, with highest occupancy occurring in coastal forests of Oregon and Washington. The 2025 field season occurred under substantial operational constraints that reduced sampling effort by approximately half relative to 2023 and 2024 because of staffing limitations affecting participating federal agencies. Despite these reductions, the NWFP-PAM framework continued to provide broad-scale, spatially representative ecological information across the NWFP area. Results highlight the growing importance of passive acoustic monitoring and machine learning approaches for long-term biodiversity monitoring under changing environmental and operational conditions.

## 1. Introduction

Northern spotted owl (*Strix occidentalis caurina*) population monitoring under the Northwest Forest Plan (NWFP) Interagency Effectiveness Monitoring Program is conducted using an integrated framework that combines passive acoustic monitoring (PAM), remote landcover sensing and mapping, and machine learning (Lesmeister and Jenkins 2022). The NWFP-PAM network provides a unified, statistically rigorous approach for assessing the status and trends of northern spotted owls, barred owls (*Strix varia*), marbled murrelets (*Brachyramphus marmoratus*), and broader forest biodiversity across federally administered lands within the NWFP area, which is defined as the geographic range of the northern spotted owl in the US. The NWFP-PAM program uses large-scale field sampling and analytical workflows to support inference on species occupancy, habitat associations, distributional change, and landscape-scale biodiversity patterns (Lesmeister and Jenkins 2022, Lesmeister et al. *in press*).

The NWFP-PAM network was developed collaboratively by an interagency group of partners and was designed to support annual surveys of more than 4,000 sampling stations distributed across approximately 24 million acres of federal forest lands. Deployments follow a randomized 2+20% sampling design (Lesmeister et al. 2021), generating spatially representative data that complement other NWFP monitoring modules focused on forest structure, habitat condition, and ecosystem change (Davis et al. 2022a, Davis et al. 2022b). From 2018–2025, the NWFP-PAM program accumulated nearly 10 million hours of acoustic recordings analyzed using the convolutional neural network PNW-Cnet, developed through the Pacific Northwest Bioacoustics Lab (Ruff et al. 2021, Ruff et al. 2023, Lesmeister et al. 2026b). These data collectively represent one of the largest long-term acoustic datasets for temperate forest ecosystems currently available.

In addition to northern spotted owl monitoring, the NWFP-PAM program reports validated detections and occupancy summaries for barred owls and marbled murrelets and provides broad-scale summaries of apparent detections for all PNW-Cnet v5 sound classes. These include birds, mammals, amphibians, anthropogenic sounds, and environmental acoustic conditions that collectively provide insight into ecological conditions across NWFP forests.

Details and summaries from previous years are available in earlier annual reports (Lesmeister et al. 2018a, Lesmeister et al. 2019, Lesmeister et al. 2022, Lesmeister et al. 2023, Lesmeister et al. 2024, Lesmeister et al. 2025).

Implementation of the 2025 field season occurred under substantial operational constraints that reduced survey capacity across participating federal agencies. Staffing limitations reduced the number of field personnel available for deployment, retrieval, logistics, and data processing, resulting in substantially lower sampling effort relative to 2023 and 2024. Continued long-term monitoring remains important for evaluating changes in northern spotted owl populations, invasive barred owl expansion, marbled murrelet habitat use, and broader biodiversity patterns under ongoing environmental change.

## 2. Study Area

We collected NWFP-PAM data within federally administered forest lands (Fig. 1):

- National Forests (*n* = 15): Deschutes, Fremont-Winema, Gifford Pinchot, Klamath, Six Rivers, Mendocino, Mount Baker-Snoqualmie, Mount Hood, Okanogan-Wenatchee, Olympic, Rogue River-Siskiyou, Shasta-Trinity, Siuslaw, Umpqua, and Willamette
- Bureau of Land Management Districts (*n* = 5): Medford, Lakeview, Coos Bay, Roseburg, and Northwest Oregon
- National Park Lands (*n* = 7): Golden Gate National Recreation Area, Muir Woods National Monument, Point Reyes National Seashore, Olympic National Park, North Cascades National Park, Mount Rainier National Park, and Redwood National Park
- Columbia River Gorge National Scenic Area that was administered by the US Forest Service Within the federal lands, we collected data within 10 historical northern spotted owl demographic study areas (Franklin et al. 2021). Nine of the study areas (OLY = Olympic Peninsula, CLE = Cle Elum, RAI = Mount Rainier National Park, COA = Oregon Coast Range, HJA = HJ Andrews Experimental Forest, TYE = Tyee, KLA = Klamath Mountains, CAS = Oregon South Cascades, NWC = Northwest California) were long-term demographic study areas for northern spotted owl monitoring under the NWFP (Franklin et al. 2021). One study area (MAR = Marin County) was included due to long-term and ongoing northern spotted owl demographic monitoring (Fig. 1). In 2025, we deployed recording units within 22 designated wilderness areas (Table 1).

**Figure 1.**
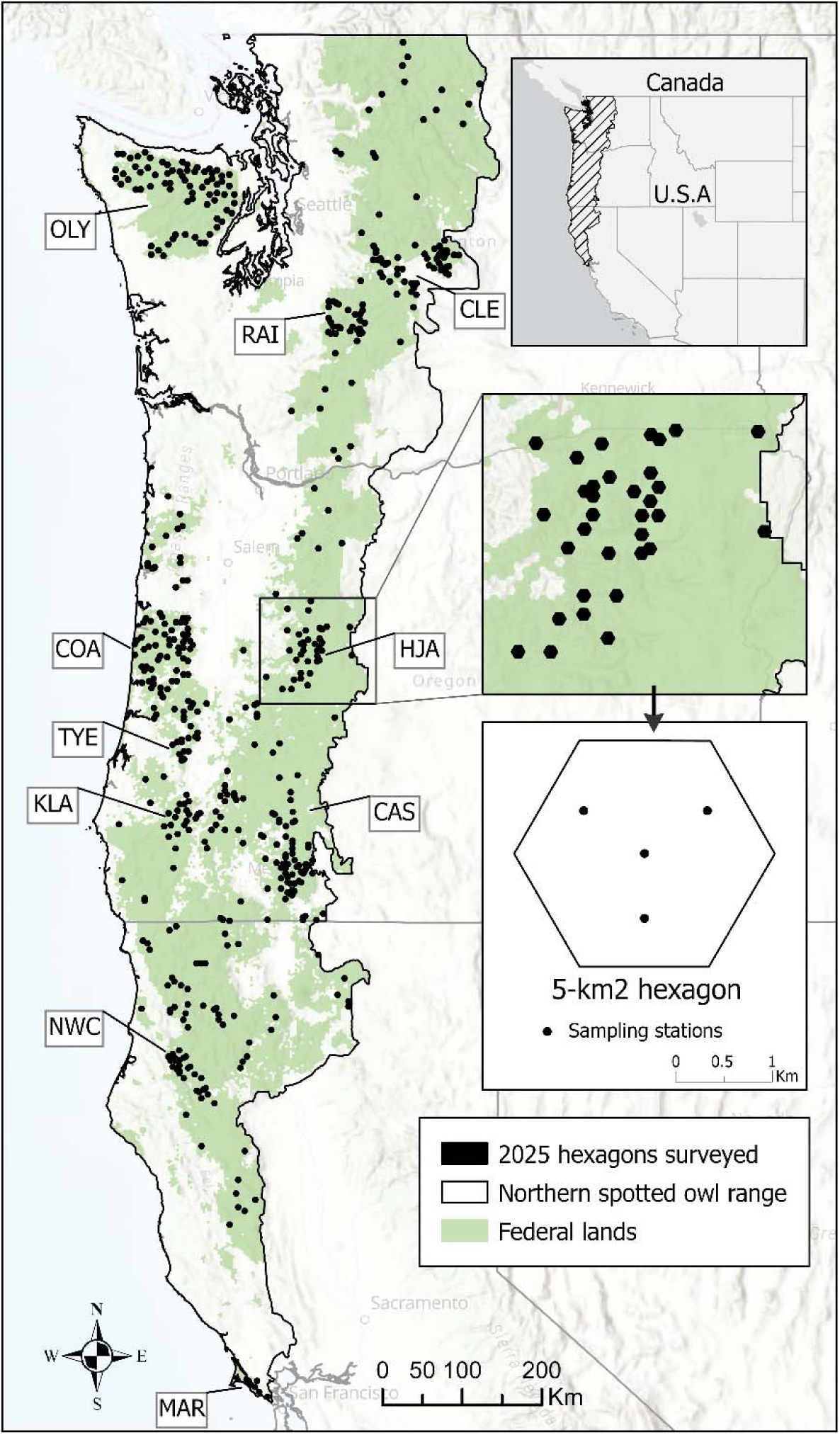
Locations of 5-km^2^ hexagons (*n* = 532) surveyed on federal lands using passive acoustic monitoring in 2025. Inset shows typical placement of acoustic sampling stations. Historical study areas sampled at 20%: COA = Oregon Coast Range, OLY = Olympic Peninsula, KLA = Klamath Mountains, CLE = Cle Elum, TYE = Tyee, HJA = HJ Andrews Experimental Forest, CAS = Oregon South Cascades, NWC = Northwest California, MAR = Marin County, and RAI = Mount Rainier National Park. WA 2%, OR 2%, and CA 2% were data collected in each state on the 2% sampling outside the 20% sampling density on historical study areas.

**Table 1.** The number of autonomous recording unit stations surveyed during 2025 in 22 designated wilderness areas administered by US Forest Service, US National Park Service, or US Bureau of Land Management.

| **Wilderness Area** | **Stations surveyed** |
| --- | --- |
| *US Forest Service* |  |
| Alpine Lakes | 8 |
| Buckhorn | 8 |
| Clackamas | 2 |
| Cummins Creek | 2 |
| Devil's Staircase | 8 |
| Drift Creek | 2 |
| Kalmiopsis | 8 |
| Lake Chelan-Sawtooth | 4 |
| Mount Hood | 4 |
| North Fork | 1 |
| Rock Creek | 1 |
| Siskiyou | 6 |
| Sky Lakes | 11 |
| The Brothers | 2 |
| Three Sisters | 5 |
| Trinity Alps | 10 |
| Yolla Bolly-Middle Eel | 4 |
| *US National Park Service* |  |
| Mount Rainier | 106 |
| Olympic | 137 |
| Phillip Burton | 7 |
| Stephen Mather | 22 |
| *US Bureau of Land Management* |  |
| Soda Mountain | 4 |
| **TOTAL** | **362** |

## 3. Methods

### Sampling design

We created a uniform layer of 5-km^2^ hexagons that covered the range of the northern spotted owl within the US (Lesmeister et al. 2021) that is available for download (USFWS 2021). This hexagon size is approximately the size of a northern spotted owl territory core area (Glenn et al. 2004, Schilling et al. 2013) and approximates the home range size reported for barred owls in the Pacific Northwest (Hamer et al. 2007, Singleton et al. 2010, Wiens et al.

2014). Within the historical demographic study areas, we randomly selected 20% of available hexagons that contained ≥50% forest capable lands and ≥25% federal ownership. Outside the historical demographic study areas, we randomly selected 2% of hexagons throughout the entire NWFP area following the same criteria for forest capable lands and federal ownership, stratified by physiographic province which describe ecologically similar regions (Thomas et al. 1993).

Forest capable lands were those areas with suitable soil type, plant association, and elevation capable of developing into forest (Davis and Lint 2005). Within each hexagon, we deployed 4 recorders (Fig 1).

We collected acoustic data at survey stations using Song Meter SM4s (*n* = 1,823) and Song Meter Minis (*n* = 272; Wildlife Acoustics, Maynard, MA), which are portable, weatherproof, and easily programmable autonomous recording units (ARUs). The SM4s had two built-in omni-directional microphones with signal-to-noise ratio of 80 decibels (dB) typical at 1 kilohertz (kHz), two SDHC/SDXC flash card slots, average of 543 h of recording battery life, and a recording bandwidth of 0.02–48.00 kHz at levels of -33.5–122.0 dB. The Song Meter Mini had the same bandwidth, signal-to-noise ratio of 78 dB, one omni-directional microphone, one SDHC/SDXC flash card slot, and 210–1040 h battery life depending on configuration. We programmed all recorders to use a sample rate of 32 kHz with one microphone. These ARUs recorded sound with equivalent sensitivity to normal range of human hearing, and their effective listening radius may be affected by external factors such as terrain, vegetation, and weather events such as wind and rain. At each sampling station within a hexagon, we mounted ARUs to small trees (15–25 cm diameter at breast height) to allow microphones to extend past the bole for unobstructed recording ability. We deployed ARUs on federal land; mid-to-upper slope positions; ≥50 m from roads, trails, and streams to reduce vandalism and excessive noise; spaced ≥500 m apart; and located ≥200 m from the edge of the hexagon. We programmed ARUs to record from 1 h before sunset to 3 h after sunset, 2 h before sunrise to 2 h after sunrise, and for the first 10 min of every hour throughout the day and night.

### Data processing

We followed a multi-step workflow that integrated PNW-Cnet v5 to efficiently process large volumes of audio data, combining automated identification and human validation (Ruff et al. 2021). PNW-Cnet v5 classifies call types for 84 species and had high precision for most sound classes (Lesmeister et al. 2026b). There were 108 sound classes with precision estimates >0.90 at the 0.95 threshold (recall ranging from 0.01–0.97). PNW-Cnet generates predictions that are interpretable as confidence scores between 0.00–1.00 for each sound class for each 12 second clip (Ruff et al. 2020). We report the number of apparent detections for all PNW-Cnet v5 classes that had at least one predicted detection, which are the number of clips with a score exceeding a classification score threshold (i.e., 0.95 for most target classes). We adjusted the number of apparent detections reported by model class precision (Lesmeister et al. 2026b). We used a high classification threshold (0.95) to maximize precision over recall for this report. These are not validated counts.

### Data validation

Apparent detections of northern spotted owl, barred owl, and marbled murrelet call classes were validated using the program Kaleidoscope (Wildlife Acoustics Inc. 2024) to examine audio and associated spectrograms. We reviewed all clips with PNW-Cnet classification scores ≥0.50 for the northern spotted owl four-note location calls and ≥0.95 for survey tones (artificial pure tones played preceding or following call-back surveys) to maximize recall. We reviewed other northern spotted owl call type classes, the marbled murrelet keer call, and the barred owl eight-note territorial call at a classification threshold of ≥0.95, reviewing as many as necessary to validate weekly station-level detection. We did not review apparent detections for marbled murrelet keer calls in hexagons outside of the NWFP murrelet management zones (Raphael et al. 2018). Northern spotted owl detections within hexagons with ≥1 confirmed spotted owl call were reviewed by at least 2 validators. During initial and secondary reviews, validators confirmed detection of species and identified sex using aspects of frequency and call duration (Dale et al. 2022).

Call-back surveys for northern spotted owls, barred owls, and, rarely, other species occurred in our study areas by biologists working on other research projects (e.g., Franklin et al. 2021, Wiens et al. 2021, Hobart et al. 2025) and project-level clearance surveys. Additionally, several regions in California and Oregon initiated or expanded barred owl control efforts in 2025, which includes call-playback during initial surveys and as lures at removal sites. These surveys broadcast recorded calls of northern spotted owls or other target species to elicit a territorial response. We distributed a recording consisting of a brief series of pure tones (1 s at 0.5, 1.5, and 1.0 kHz) for *Strix* call-back surveyors to voluntarily play at the same volume directly before or after northern spotted owl call-back surveys (USFWS 2021). We removed any validated detections of the surveyed species if there was a reported or suspected call-back survey in the hexagon on the same night. These surveyors were included in the survey request and we removed call detections overlapping removal surveys as well.

We report the counts of validated detections of male and female northern spotted owl four-note location calls and survey tones and the proportion of surveyed hexagons with validated detections for northern spotted owls, marbled murrelets, and barred owls by historical study area (20% sample) and the 2% sample in each state.

### Background noise analysis

Background noise has consistently been found to be an important predictor of detection probabilities in species occurrence models (Duchac et al. 2020, Duchac et al. 2021, Rugg et al. 2025). Therefore, we used the sound pressure level analysis feature in Kaleidoscope Pro to quantify relative background noise levels at each sampling station which are available upon request but not reported here.

## 4. Results

In 2025, we surveyed stations in 532 hexagons with 1.37 million h of recordings, representing roughly half of sites sampled in 2023 and 2024 (Table 2). Reductions were well dispersed spatially, with most study areas sampled at 40–60% of previous completed; MAR was fully sampled, 90% of samples were attained in NWC and 96% in RAI (Table 2). Deployments took place from March 1–July 31, 2025. We deployed ARUs at 362 sampling stations in wilderness areas administered by US National Park Service (*n* = 272), US Forest Service (*n* = 86) or US Bureau of Land Management (*n* = 4; Table 1). Each year we experience some degree of data loss by various sources such as wildfire, theft, animal damage, water intrusion, and software or hardware failure. Additionally, limited staff and fire closures pushed retrievals later than ordinary resulting in three hexagons left uncollected until the following year.

**Table 2.** Passive acoustic monitoring effort during 2018–2025 within in the Northwest Forest Plan Area, summarized by historical study areas (i.e., 20% sampling) and 2% sampling (outside 20% sample density) by state. Historical study areas: COA = Oregon Coast Range, OLY = Olympic Peninsula, KLA = Klamath, CLE = Cle Elum, TYE = Tyee, HJA = HJ Andrews Experimental Forest, CAS = Oregon South Cascades, NWC = Northwest California, MAR = Marin County, and RAI = Mount Rainier National Park.

| Area | Number of hexagons |  |  |  |  |  |  |  | Number of stations |  |  |  |  |  |  |  |
| --- | --- | --- | --- | --- | --- | --- | --- | --- | --- | --- | --- | --- | --- | --- | --- | --- |
|  | 2018 | 2019 | 2020 | 2021 | 2022 | 2023 | 2024 | 2025 | 2018<sup>a</sup> | 2019 | 2020 | 2021 | 2022 | 2023 | 2024 | 2025 |
| COA | 120 | 106 | 120 | 120 | 117 | 120 | 120 | 59 | 578 | 413 | 471 | 475 | 466 | 476 | 479 | 231 |
| OLY | 88 | 120 | 119 | 119 | 120 | 120 | 118 | 73 | 436 | 472 | 464 | 472 | 474 | 479 | 472 | 285 |
| KLA |  | 63 | 73 | 73 | 73 | 73 | 73 | 31 |  | 244 | 290 | 276 | 292 | 292 | 291 | 123 |
| CLE |  |  | 69 | 75 | 75 | 75 | 75 | 43 |  |  | 267 | 298 | 299 | 300 | 300 | 171 |
| TYE |  |  |  | 40 | 40 | 40 | 40 | 22 |  |  |  | 155 | 160 | 160 | 160 | 83 |
| HJA |  |  |  | 70 | 70 | 70 | 69 | 32 |  |  |  | 279 | 280 | 278 | 273 | 126 |
| CAS |  |  |  | 98 | 98 | 98 | 98 | 46 |  |  |  | 380 | 391 | 391 | 389 | 184 |
| NWC |  |  |  | 30 | 30 | 30 | 30 | 27 |  |  |  | 120 | 119 | 118 | 117 | 103 |
| MAR |  |  |  | 7 | 7 | 7 | 7 | 7 |  |  |  | 26 | 27 | 27 | 28 | 28 |
| RAI |  |  |  | 11 | 24 | 32 | 28 | 27 |  |  |  | 43 | 96 | 128 | 111 | 106 |
| WA 2% |  |  |  |  | 23<sup>b</sup> | 90 | 81 | 35 |  |  |  |  | 91<sup>b</sup> | 355 | 323 | 139 |
| OR 2% |  |  |  |  | 13<sup>b</sup> | 178 | 186 | 79 |  |  |  |  | 52<sup>b</sup> | 709 | 732 | 315 |
| CA 2% |  |  |  |  |  | 76 | 95 | 51 |  |  |  |  |  | 299 | 377 | 201 |
| Total | 208 | 289 | 381 | 643 | 690 | 1,009 | 1,020 | 532 | 1,014 | 716 | 464 | 2,524 | 2,747 | 4,012 | 4,052 | 2,095 |

<sup>a</sup> During 2018 survey design was five stations per hexagon.
<sup>b</sup> In 2022, Gifford Pinchot National Forest in Washington and Umpqua National Forest in Oregon were sampled at 2%.

In 2025, we received broadcast survey occurrence records for barred owl and northern spotted owl call-back surveys within or adjacent to our sampled hexagons. *Strix* owl call-back surveys were reported in 17% of our hexagons (*n* = 92) and we found the artificial tone in 77 hexagons (range of 1–47 clips per hexagon). We removed some northern spotted owl calls in 95 hexagons due to reported or suspected overlapping call-back survey records. The KLA area had the highest number of survey tone detections (43% of detected tones). No call-back surveys were reported in CLE, MAR, OLY, or RAI and no survey tones were found in those areas or CA 2% areas (Table 4).

**Table 3.** Thousands of hours of passive acoustic monitoring data collected during 2018–2025 in each study area and processed for automated species identification with PNW-Cnet. Historical study areas: COA = Oregon Coast Range, OLY = Olympic Peninsula, KLA = Klamath, CLE = Cle Elum, TYE = Tyee, HJA = HJ Andrews Experimental Forest, CAS = Oregon South Cascades, NWC = Northwest California, MAR = Marin County, and RAI = Mount Rainier National Park. WA 2%, OR 2%, and CA 2% were data collected in each state on the 2% sampling outside the 20% sampling density on historical study areas.

| Area | 2018<sup>a</sup> | 2019 | 2020 | 2021 | 2022 | 2023 | 2024 | 2025 |
| --- | --- | --- | --- | --- | --- | --- | --- | --- |
| COA | 197 | 155 | 187 | 217 | 258 | 259 | 267 | 155 |
| OLY | 147 | 184 | 219 | 215 | 264 | 271 | 264 | 172 |
| KLA |  | 92 | 130 | 135 | 162 | 171 | 171 | 83 |
| CLE |  |  | 104 | 153 | 152 | 168 | 160 | 106 |
| TYE |  |  |  | 78 | 92 | 88 | 89 | 59 |
| HJA |  |  |  | 131 | 150 | 149 | 152 | 84 |
| CAS |  |  |  | 178 | 188 | 195 | 199 | 126 |
| NWC |  |  |  | 48 | 54 | 62 | 62 | 67 |
| MAR |  |  |  | 11 | 10 | 15 | 15 | 19 |
| RAI |  |  |  | 11 | 51 | 71 | 56 | 56 |
| WA 2% |  |  |  |  | 44<sup>b</sup> | 192 | 186 | 88 |
| OR 2% |  |  |  |  | 30<sup>b</sup> | 385 | 420 | 218 |
| CA 2% |  |  |  |  |  | 151 | 204 | 137 |
| Total | 344 | 431 | 640 | 1,177 | 1,456 | 2,179 | 2,244 | 1,370 |

<sup>a</sup> During 2018 survey design was five stations per hexagon.
<sup>b</sup> In 2022, Gifford Pinchot National Forest in Washington and Umpqua National Forest in Oregon were sampled at 2%.

**Table 4.**
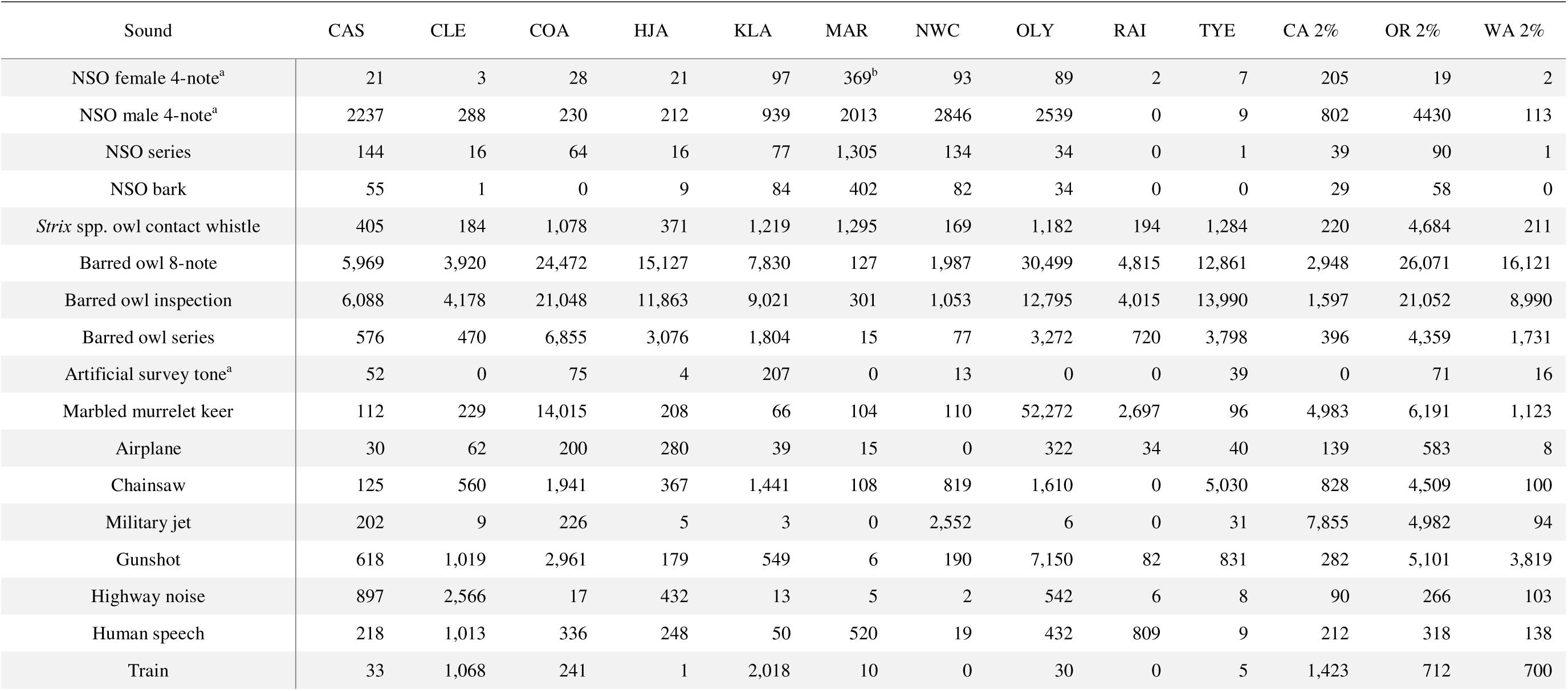
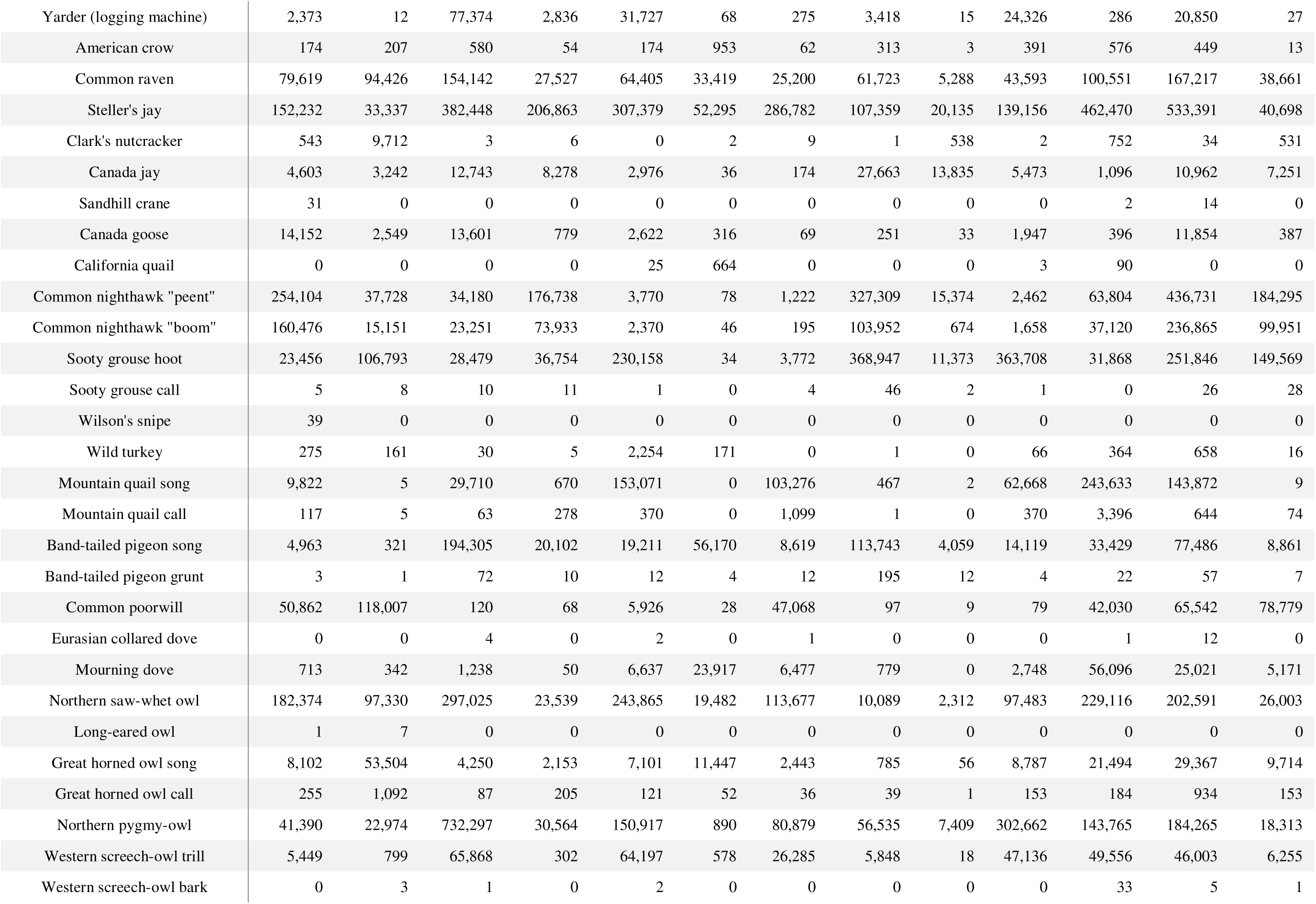
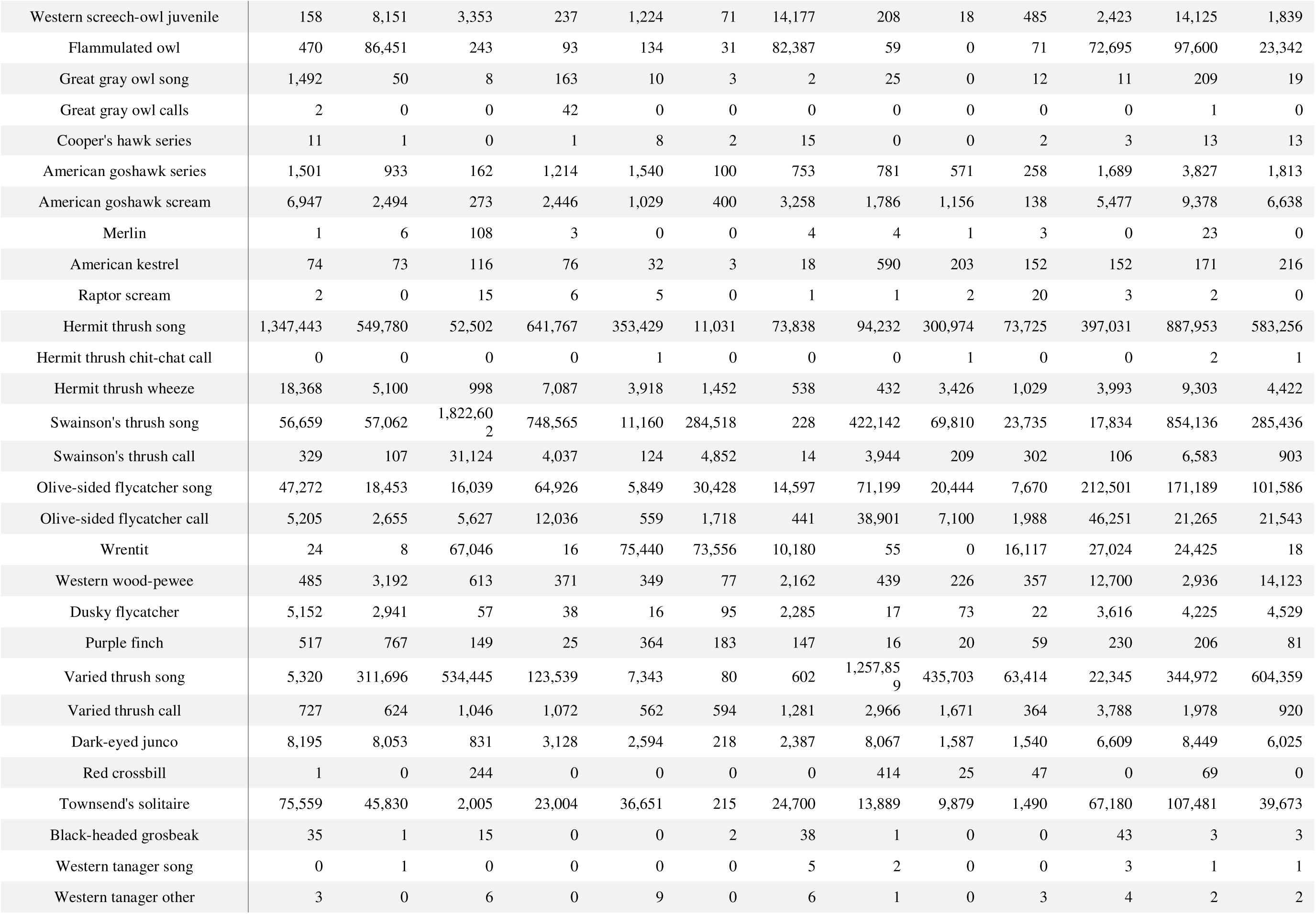
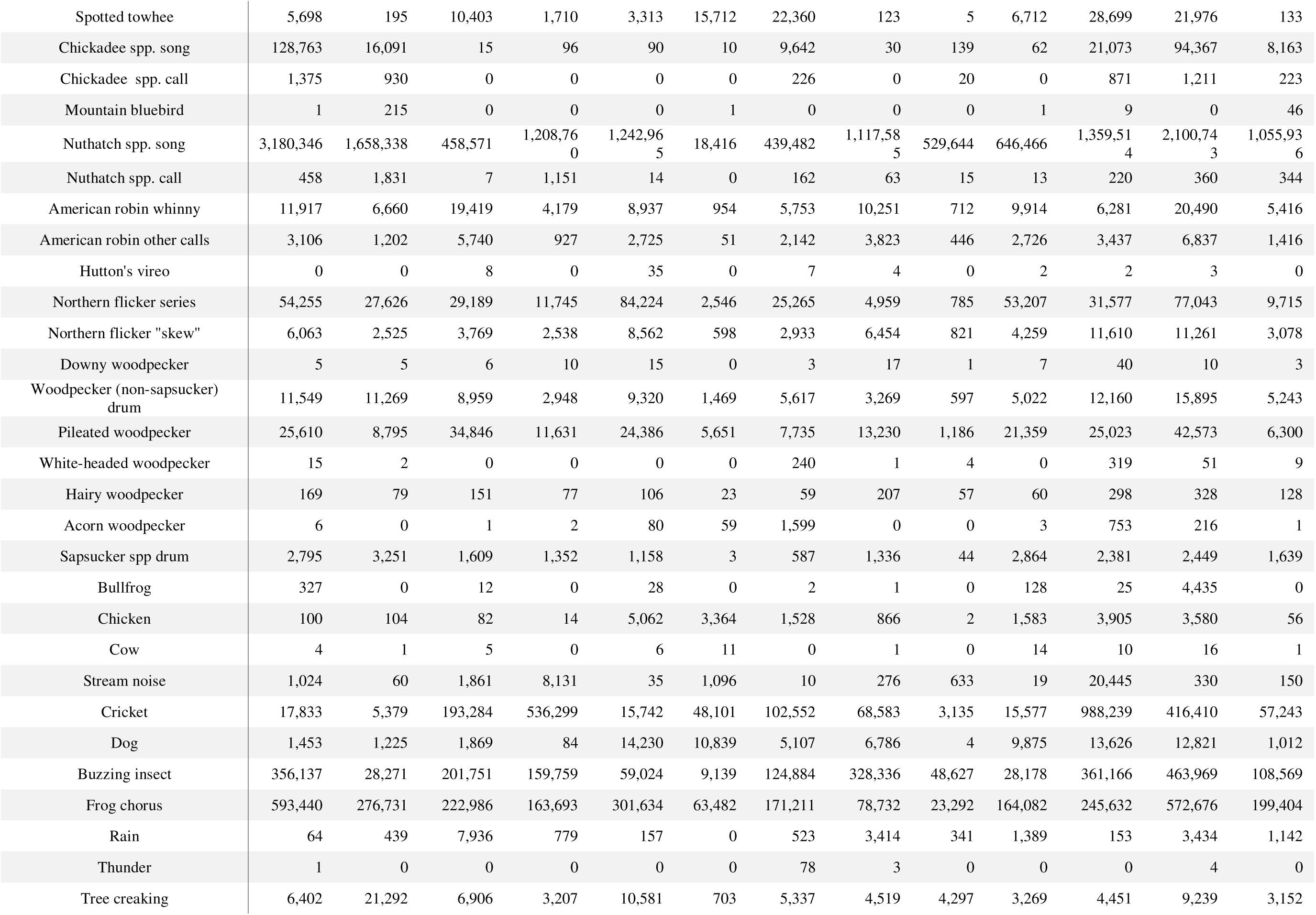
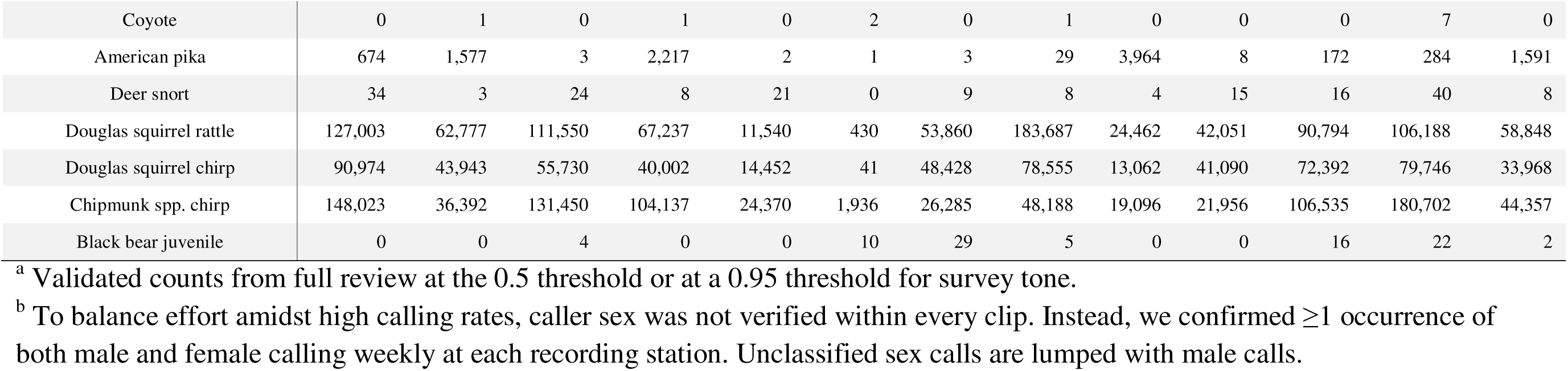
Predicted number of detections of PNW-Cnet v5 sound classes from passive acoustic monitoring data by study area collected in 2025. Predicted detections for each sound class were calculated as the number of 12-second clips in the audio dataset to which the PNW-Cnet v5 assigned a score exceeding 0.95 for that class, multiplied by the precision estimate. Validated call counts are given for the primary northern spotted owl call (NSO; after call-back survey removal) and survey tone call classes which were fully reviewed at 0.5 and 0.95 thresholds, respectively. Validated call counts are given for barred owl 8-note and marbled murrelet keer calls above 0.95 thresholds. Classes with zero apparent detections not shown (e.g., gray wolf). Historical NSO study areas (20% sampling): CAS = Oregon South Cascades, OR; CLE = Cle Elum, WA; COA = Oregon Coast Range, OR; HJA = HJ Andrews Experimental Forest, OR; KLA = Klamath Mountains, OR and CA; MAR = Marin County, CA; NWC = Northwest California, CA; OLY = Olympic Peninsula, WA; RAI = Mount Rainier National Park, WA; and TYE = Tyee, OR. 2% random sample of federal lands in Washington (WA 2%), Oregon (OR 2%), and California (CA 2%).

### Focal species detections

In 2025, we observed northern spotted owls in 28% of sampled hexagons range-wide (16% in WA, 25% OR, 60% CA), including detections in all 20% and 2% areas (Fig. 2). As in previous years, we found mostly (95%) male spotted owls compared to female spotted owls (Table 4). California study areas (MAR, NWC) had the highest proportion of hexagons with spotted owl detections and MAR had the most female detections despite having the fewest sites (Table 5). Study areas in the Washington Cascades (CLE, RAI, WA 2%), and the COA study area had some of the lowest proportion of hexagons with detections (Table 5). The proportion of hexagons with detections fluctuated in most study areas with multiple years of surveys (Table 5). The TYE study area declined from 38% (15 hexagons) to 18% (4 hexagons) from 2021–2025.

**Figure 2.**
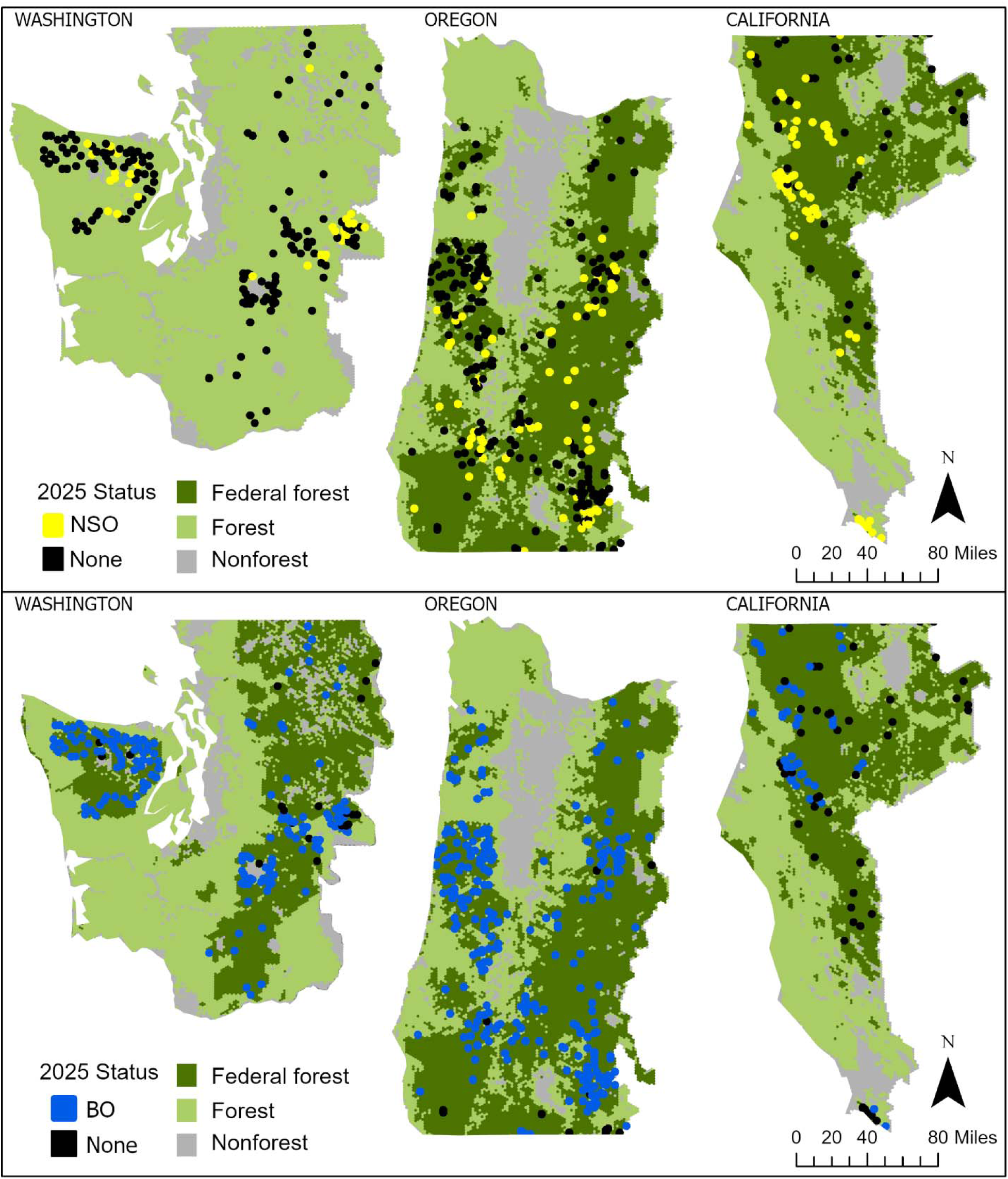
Locations of 5-km^2^ hexagons with validated northern spotted owl (NSO: *n* =149) and barred owl (BO; *n* = 455) detections in 2024 acoustic monitoring.

**Table 5.** Proportion of monitored hexagons with validated detections of northern spotted owl for years that surveys were conducted (2018–2024) within the Northwest Forest Plan Area. Historical study areas sampled at 20%: COA = Oregon Coast Range, OLY = Olympic Peninsula, KLA = Klamath Mountains, CLE = Cle Elum, TYE = Tyee, HJA = HJ Andrews Experimental Forest, CAS = Oregon South Cascades, NWC = Northwest California, MAR = Marin County, and RAI = Mount Rainier National Park. WA 2%, OR 2%, and CA 2% were data collected in each state on the 2% sampling outside the 20% sampling density on historical study areas.

| Area | 2018 | 2019 | 2020 | 2021 | 2022 | 2023 | 2024 | 2025 |
| --- | --- | --- | --- | --- | --- | --- | --- | --- |
| COA | 0.16 | 0.13 | 0.09 | 0.15 | 0.09 | 0.11 | 0.10 | 0.12 |
| OLY | 0.16 | 0.17 | 0.24 | 0.19 | 0.27 | 0.25 | 0.14 | 0.18 |
| KLA |  | 0.43 | 0.53 | 0.49 | 0.51 | 0.37 | 0.23 | 0.39 |
| CLE |  |  | 0.19 | 0.19 | 0.11 | 0.20 | 0.13 | 0.26 |
| TYE |  |  |  | 0.38 | 0.28 | 0.15 | 0.05 | 0.18 |
| HJA |  |  |  | 0.39 | 0.39 | 0.43 | 0.35 | 0.34 |
| CAS |  |  |  | 0.35 | 0.31 | 0.23 | 0.31 | 0.28 |
| NWC |  |  |  | 0.77 | 0.73 | 0.80 | 0.60 | 0.96 |
| MAR |  |  |  | 1.00 | 1.00 | 1.00 | 1.00 | 1.00 |
| RAI |  |  |  | 0.09 | 0.04 | 0.06 | 0.07 | 0.04 |
| WA 2% |  |  |  |  | 0.00<sup>a</sup> | 0.09 | 0.07 | 0.11 |
| OR 2% |  |  |  |  | 0.54<sup>a</sup> | 0.24 | 0.17 | 0.24 |
| CA 2% |  |  |  |  |  | 0.37 | 0.38 | 0.35 |

<sup>a</sup> In 2022, Gifford Pinchot National Forest in Washington and Umpqua National Forest in Oregon were sampled at 2%.

While KLA has declined since 2022 from 51% (37 hexagons) to 39% (12 hexagons) in 2025 (Table 5). We did not detect spotted owls in Columbia River Gorge NSA, Deschutes National Forest, Gifford Pinchot National Forest, Mt. Baker-Snoqualmie National Forest, Mt. Hood National Forest, or the Lakeview District BLM. We detected male spotted owl calls at one hexagon in North Cascades National Park.

We detected barred owl calling in 86% of surveyed hexagons (Table 6; Figure 2). In Washington, we detected barred owls in 89% of surveyed hexagons (*n* = 1,580). In Oregon, we detected barred owls in 96% of hexagons (*n* = 257; Figure 2). We found barred owls in 47% of California hexagons (*n* = 40) and CA study areas had lower proportions of hexagons and numbers of classified barred owl 8-note calls compared to sites in Oregon and Washington (Table 4, Table 6). Mendocino National Forest in California was the only surveyed federal management unit with no barred owl calls detected.

**Table 6.** Proportion of monitoring hexagons with validated detections of barred owls for years that surveys were conducted (2018–2025) within the Northwest Forest Plan Area. Historical study areas sampled at 20%: COA = Oregon Coast Range, OLY = Olympic Peninsula, KLA = Klamath Mountains, CLE = Cle Elum, TYE = Tyee, HJA = HJ Andrews Experimental Forest, CAS = Oregon South Cascades, NWC = Northwest California, MAR = Marin County, and RAI = Mount Rainier National Park. WA 2%, OR 2%, and CA 2% were data collected in each state on the 2% sampling outside the 20% sampling density on historical study areas.

| Area | 2018 | 2019 | 2020 | 2021 | 2022<sup>a</sup> | 2023 | 2024 | 2025 |
| --- | --- | --- | --- | --- | --- | --- | --- | --- |
| COA | 0.99 | 0.99 | 1.00 | 0.99 | 1.00 | 0.99 | 1.00 | 1.00 |
| OLY | 0.92 | 0.93 | 0.93 | 0.90 | 0.94 | 0.93 | 0.95 | 0.96 |
| KLA |  | 0.86 | 0.95 | 0.95 | 1.00 | 0.97 | 0.92 | 0.97 |
| CLE |  |  | 0.77 | 0.65 | 0.79 | 0.80 | 0.73 | 0.79 |
| TYE |  |  |  | 1.00 | 1.00 | 1.00 | 1.00 | 1.00 |
| HJA |  |  |  | 0.99 | 0.99 | 0.97 | 0.96 | 0.97 |
| CAS |  |  |  | 0.95 | 0.96 | 0.94 | 0.93 | 0.98 |
| NWC |  |  |  | 0.73 | 0.73 | 0.67 | 0.67 | 0.63 |
| MAR |  |  |  | 0.29 | 0.86 | 0.14 | 0.29 | 0.29 |
| RAI |  |  |  | 0.82 | 0.88 | 0.94 | 0.96 | 0.96 |
| WA 2% |  |  |  |  | 1.00<sup>b</sup> | 0.93 | 0.89 | 0.80 |
| OR 2% |  |  |  |  | 1.00<sup>b</sup> | 0.84 | 0.83 | 0.89 |
| CA 2% |  |  |  |  |  | 0.37 | 0.33 | 0.41 |

<sup>a</sup> In 2022, we didn't fully validate barred owl classes, adjusted PNW-Cnet model predicted proportions are reported.
<sup>b</sup> In 2022, only Gifford Pinchot National Forest in Washington and Umpqua National Forest in Oregon were sampled at 2%.

Marbled murrelet keer calls were identified in 50% of hexagons within NWFP marbled murrelet management zones (Figure 3). Marbled murrelets were commonly detected (78–96% of hexagons) in the two most coastal 20% study areas (COA, OLY) (Table 7; Figure 3). A higher proportion of the OR 2% sampled areas had murrelet detections compared to WA 2% and CA 2%, where sampling was further from coastal forest (Figure 3; Table 7).

**Figure 3.**
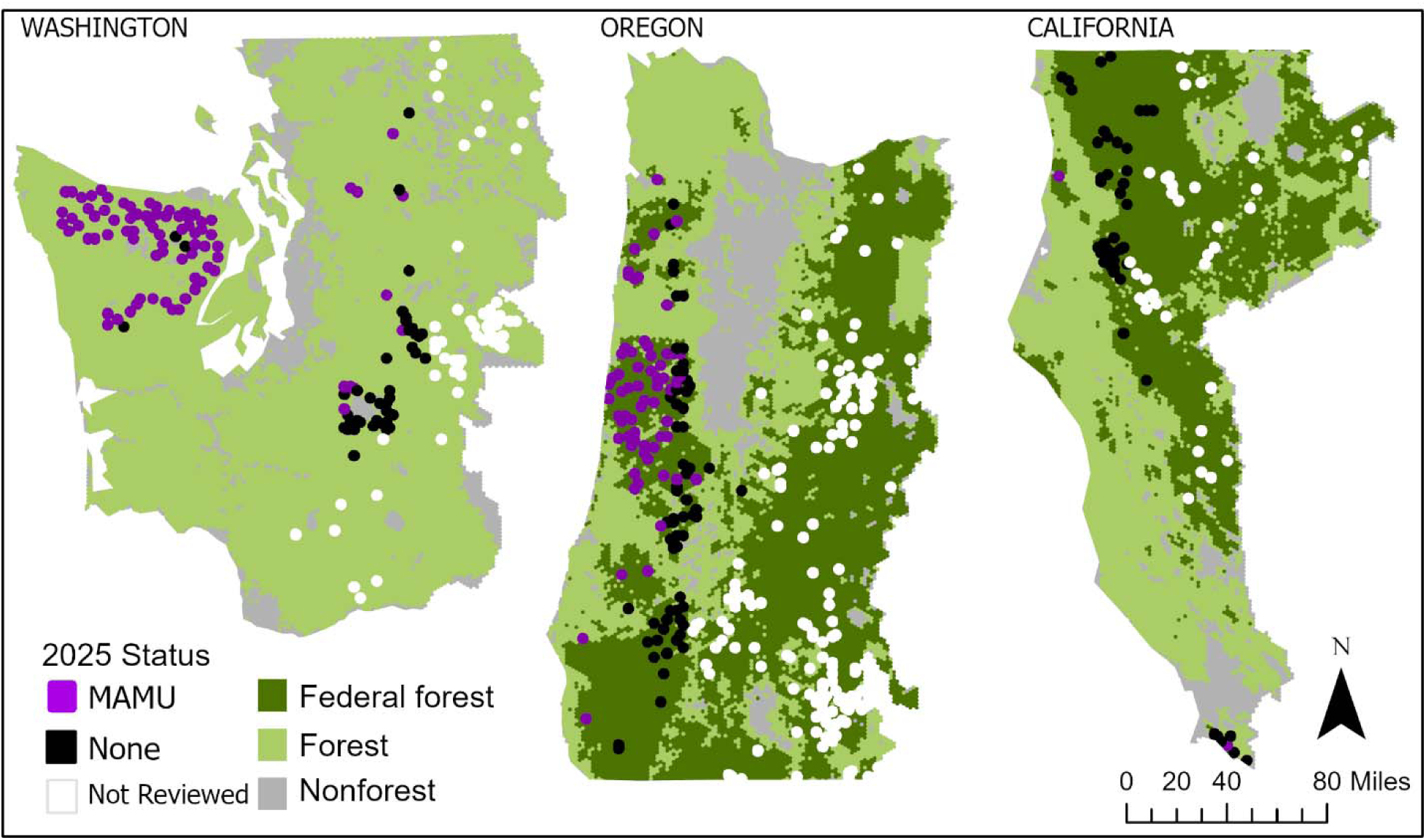
Locations of 5-km^2^ hexagons with validated marbled murrelet (MAMU; *n* = 146) detections in 2024. Only hexagons within the marbled murrelet Northwest Forest Plan management zones were reviewed (*n* = 294).

**Table 7.** Proportion of monitoring hexagons with validated detections of marbled murrelet for years that surveys were conducted (2018–2025) within the Northwest Forest Plan Area. Historical study areas sampled at 20%: COA = Oregon Coast Range, OLY = Olympic Peninsula, KLA = Klamath Mountains, CLE = Cle Elum, TYE = Tyee, HJA = HJ Andrews Experimental Forest, CAS = Oregon South Cascades, NWC = Northwest California, MAR = Marin County, and RAI = Mount Rainier National Park. WA 2%, OR 2%, and CA 2% were data collected in each state on the 2% sampling outside the 20% sampling density on historical study areas.

| Area | 2018 | 2019 | 2020 | 2021 | 2022 | 2023 | 2024<sup>c</sup> | 2025<sup>c</sup> |
| --- | --- | --- | --- | --- | --- | --- | --- | --- |
| COA | 0.83 | 0.75 | 0.79 | 0.82 | 0.86 | 0.91 | 0.87 | 0.78 |
| OLY | 0.78 | 0.85 | 0.84 | 0.87 | 0.85 | 0.92 | 0.97 | 0.96 |
| KLA |  | 0.00 | 0.00 | 0.01 | 0.00 | 0.01 | 0.01 | 0.00 |
| CLE |  |  | 0.00 | 0.00 | 0.00 | 0.00 | 0.00 | 0.02 |
| TYE |  |  |  | 0.15 | 0.10 | 0.05 | 0.07 | 0.14 |
| HJA |  |  |  | 0.00 | 0.00 | 0.00 | 0.00<sup>b</sup> | 0.00<sup>b</sup> |
| CAS |  |  |  | 0.00 | 0.00 | 0.00 | 0.00<sup>b</sup> | 0.00<sup>b</sup> |
| NWC |  |  |  | 0.00 | 0.00 | 0.00 | 0.00 | 0.00 |
| MAR |  |  |  | 0.00 | 0.00 | 0.00 | 0.00 | 0.14 |
| RAI |  |  |  | 0.09 | 0.13 | 0.13 | 0.14 | 0.15 |
| WA 2% |  |  |  |  | 0.00 | 0.13 | 0.05 | 0.14 |
| OR 2% |  |  |  |  | 0.08 | 0.20 | 0.19 | 0.19 |
| CA 2% |  |  |  |  |  | 0.01 | 0.02 | 0.02 |

<sup>a</sup> In 2022, only Gifford Pinchot National Forest in Washington and Umpqua National Forest in Oregon were sampled at 2%.
<sup>b</sup> No sites reviewed for marbled murrelet.
<sup>c</sup> Only hexagons within the marbled murrelet Northwest Forest Plan conservation zones were reviewed in 2024 & 2025.

Of the other owl species, northern saw-whet owls (*Aegolius acadicus*) and northern pygmy owls (*Glaucidium californicum)* had the greatest number of predicted vocalizations (i.e., not validated) throughout our study (Table 4). We found >1,000 apparent detections of great gray owls in CAS and >100 in HJA, both areas that we have previously confirmed presence (Table 4). Flammulated owls (*Psiloscops flammeolus*) were most frequently predicted (>20,000–97,000) in CLE, NWC, and the 2% sample areas in all three states (Table 4). Great horned owl *(Bubo virginianus*) and western screech-owl (*Megascops kennicottii*) calls were predicted in all sampling areas (range: 56–53,504 and 18–65,868, respectively). We found a few apparent long-eared owl (*Asio otus*) calls in CAS and CLE (Table 4).

The highest proportions of American goshawk (*Accipiter gentilis*) predicted detections were in the 2% areas and in the CAS study area (10–23% of predicted calls study-wide).

American pika (*Ochotona princeps*) were commonly detected (>1,000) in CLE, HJA, RAI and WA 2%. Band-tailed pigeon (*Patagioenas fasciata*) call classes were predicted in all areas with as few as 321 in CLE up to 194,305 apparent detections in COA (Table 4). Sooty grouse (*Dendragapus fuliginosus*) call classes were predicted most densely (>300,000) in OLY and TYE.

We did not find apparent detections above the 0.95 threshold for sharp-shinned hawk (*Accipiter striatus*), ruffed grouse (*Bonasa umbellus*), wolf (*Canis lupus*), Wilson’s warbler (*Cardellina pusilla*), elk (*Cervus canadensis*), Pacific-slope flycatcher, bald eagle (*Haliaeetus leucocephalus*), evening grosbeak (*Hesperiphona vespertina*), car horn, orange-crowned warbler (*Leiothlypis celata*), Osprey (*Pandion haliaetus*), chipping sparrow (*Spizella passerine*), pine siskin (*Spinus pinus*), Williamson’s sapsucker (*Sphyrapicus thyroideus*), northern house wren (*Troglodytes aedon*), wildcat (*Puma concolor* and *Lynx rufus*), or white-crowned sparrow (*Zonotrichia leucophrys*). Each of these species were new classes for PNW-Cnet.v5 with small training datasets (Lesmeister et al. 2026b), so the lack of apparent detections is likely from poor model performance rather than absence from areas surveyed. For many of these species the recall in PNW-Cnet v5 may have been too low for any true positives to reach the 0.95 classification score threshold used in this report.

## 5. Discussion

We report PAM results collected during 2025 within the NWFP area, representing the eighth consecutive year of large-scale implementation and the third year of full implementation of the range-wide 2+20% sampling framework (Lesmeister et al. 2021). Although the NWFP-PAM program was originally developed to support northern spotted owl effectiveness monitoring, the program now provides one of the most spatially extensive biodiversity monitoring datasets available for federal forest lands in the Pacific Northwest. The 2025 field season continued to demonstrate the value of NWFP-PAM for simultaneously monitoring northern spotted owls, invasive barred owls, marbled murrelets, and broader vertebrate biodiversity using a unified survey framework. Increasingly, NWFP-PAM datasets are supporting ecological inference and model development beyond species occupancy alone, including phenology, soundscape ecology, community interactions, and sensory ecology applications enabled through advances in artificial intelligence and automated acoustic classification (Weldy et al. 2024, Wiens et al. 2025, Clapp et al. 2026, Habib et al. 2026, Weldy et al. 2026).

### Northern spotted owl

Northern spotted owl occupancy continued to exhibit strong geographic structure across the NWFP area, with higher rates of detection occurring in southern regions and persistently lower occupancy in the Washington Cascades and portions of the Oregon Coast Range. Across the 2021–2025 monitoring period, occupancy patterns generally paralleled long-term demographic trends documented by Franklin et al. (2021) and (Dugger et al. *in review*), including continued evidence of contraction in portions of Washington and northern Oregon where barred owl occupancy is now extensive.

In 2025, northern spotted owls were detected in 28% of sampled hexagons range-wide, including 16% of hexagons in Washington, 25% in Oregon, and 60% in California. California study areas continued to have the highest occupancy rates, particularly in Marin County and northwest California, where northern spotted owl detections remained relatively widespread and barred owl occupancy remained comparatively lower than in much of Oregon and Washington. These results remain consistent with analyses linking northern spotted owl persistence to regions with reduced barred owl occupancy and higher concentrations of mature forest habitat (Wiens et al. 2021, Jenkins et al. 2025, Wiens et al. 2025, Lesmeister et al. *in press*).

For the first time since range-wide deployment began in 2023, northern spotted owl detections were confirmed within North Cascades National Park in the Washington Cascades sampling region. Although this detection is encouraging, it should not be interpreted as evidence of regional population recovery. Rather, the observation may reflect dispersal by isolated individuals into landscapes where occupancy has become extremely low following decades of barred owl expansion. The continued rarity of detections throughout much of the Washington Cascades reinforces concerns regarding long-term persistence in the northern portion of the range (Lesmeister et al. *in press*).

The southern 20% study areas Klamath Mountains and Northwest California showed increases or stabilization in spotted owl occupancy relative to recent years. Concurrently, barred owl management actions expanded in northwestern California and southern Oregon during 2024–2025 (Hobart et al. 2025). Although causal relationships cannot yet be inferred from these monitoring data alone, these patterns are consistent with previous experimental evidence demonstrating demographic benefits of barred owl removal for northern spotted owls (Wiens et al. 2021, Wiens et al. 2025). Continued monitoring will be important for evaluating whether localized increases in occupancy persist through time in areas with active barred owl management.

Recent work refining occupancy inference from PAM has also highlighted the importance of interpreting spotted owl vocal activity within an ecological and behavioral context (Rugg et al. 2025). Vocal detections likely reflect varying combinations of territorial behavior, pair occupancy, and landscape use rather than simple presence or absence alone. As long-term datasets continue to expand, integration of behavioral ecology with PAM frameworks may improve interpretation of occupancy dynamics and strengthen inference regarding northern spotted owl population status and habitat use.

Collectively, these findings indicate that northern spotted owl persistence may increasingly depend on a combination of remaining high-quality habitat, localized refugia from barred owl dominance, and active management interventions (Lesmeister et al. 2018b, Wiens et al. 2021, Dumbacher and Franklin 2024). The NWFP-PAM framework provides an important mechanism for tracking these patterns consistently across large spatial scales and identifying areas where conservation actions may have the greatest long-term value.

### Barred owl

Barred owl detections remained widespread throughout most of the NWFP area in 2025, with validated detections occurring in 86% of sampled hexagons. Occupancy remained especially high throughout western Oregon and Washington, where many study areas now consistently exhibit barred owl detections in >90% of sampled hexagons. These patterns continue to support previous analyses demonstrating broad landscape dominance of barred owls across large portions of the northern spotted owl range (Jenkins et al. 2025).

The consistency in barred owl distribution patterns reinforces the suggestion that the NWFP-PAM framework is reliable for tracking invasive species populations across broad geographic regions. Areas with the highest rates of barred owl occupancy continued to overlap with regions where northern spotted owl occupancy has declined most substantially, particularly in the Oregon Coast Range and portions of the Washington Cascades. In contrast, portions of California continued to exhibit comparatively lower barred owl occupancy and higher northern spotted owl occupancy.

The consistency of these patterns across years provides an important baseline for evaluating outcomes of ongoing barred owl management efforts. Recent work examining owl community responses following barred owl removal demonstrated that PAM can effectively quantify community-level ecological responses to invasive species management, including changes in occupancy dynamics among sympatric owl species (Wiens et al. 2025). The NWFP-PAM network is increasingly serving as a complementary monitoring framework by providing broad-scale occupancy information across large landscapes. As barred owl management moves forward, continued coordination among research and management partners will be important for evaluating long-term responses of both species to changing management strategies and landscape conditions.

### Marbled murrelet

Marbled murrelet monitoring continued to demonstrate the value of NWFP-PAM for assessing occupancy across large coastal forest landscapes. Region-wide deployments now encompass more than 200 coastal sites extending from northern California to Washington’s Olympic Peninsula, providing one of the most extensive acoustic monitoring datasets currently available for inland murrelet habitat use.

Validated detections remained concentrated within coastal forests of the Olympic Peninsula and Oregon Coast Range, where occupancy rates remained consistently high across years. In 2025, marbled murrelets were detected in 50% of reviewed hexagons within NWFP marbled murrelet management zones, with the highest rates occurring in the COA and OLY study areas. Patterns remained strongly associated with mature coastal conifer forests and low-fragmentation landscapes, consistent with established nesting habitat relationships (Raphael et al. 2018, Lorenz et al. 2021).

Results from the NWFP-PAM framework continue to support recent findings demonstrating that the network can provide a cost-effective alternative to traditional inland audio-visual surveys while maintaining high detection probabilities (Duarte et al. 2024, Thomas et al. 2025). The ability to deploy ARUs across remote coastal terrain for extended sampling periods allows for substantially broader spatial coverage while reducing personnel requirements and survey costs.

Together, these findings position the NWFP-PAM framework as an increasingly important component of long-term marbled murrelet monitoring under the NWFP. Continued integration of PAM into murrelet monitoring programs may improve the ability to track changes in inland habitat use across broad spatial and temporal scales.

### Biodiversity

Beyond focal species monitoring, the 2025 NWFP-PAM dataset continued to demonstrate the broader ecological value of large-scale PAM networks. PNW-Cnet v5 generated apparent detections for a wide range of birds, mammals, amphibians, anthropogenic sounds, and environmental acoustic conditions, providing a broad snapshot of ecological and soundscape conditions across Pacific Northwest forests. Increasingly, these datasets are functioning not only as wildlife monitoring records, but also as long-term ecological archives capable of supporting retrospective analyses and new forms of biodiversity inference as analytical approaches continue to evolve.

Many species exhibited strong geographic structure consistent with known habitat associations and distributions. For example, flammulated owls were most detected in drier forest landscapes east of the Cascade crest, while marbled murrelets remained concentrated in coastal forests. American pika detections were associated primarily with higher elevation landscapes, and mountain quail detections were concentrated in southwestern Oregon and northwestern California. Broad distributions of species such as varied thrush (*Ixoreus naevius*), hermit thrush (*Catharus guttatus*), northern pygmy-owl (*Glaucidium californicum*), and Douglas squirrel (*Tamiasciurus douglasii*) further illustrate the ability of the NWFP-PAM framework to capture biologically meaningful landscape-scale patterns across diverse taxa. Several species generated apparent detections outside their currently documented geographic ranges. Because these observations may reflect either true range expansions or false positives associated with limited training data or lower model precision for some newly added classes, further validation of these detections is needed before they can be interpreted as confirmed occurrences.

The 2025 field season also highlighted the continued value of bioacoustic bycatch, where monitoring originally designed for focal species simultaneously generates information relevant to broader biodiversity objectives. These datasets increasingly support analyses related to wildfire effects, habitat change, species distributions, ecological indicators, and soundscape ecology. For example, NWFP-PAM data was effectively used to quantify broad-scale occupancy patterns of pileated woodpeckers (*Dryocopus pileatus*) across the Pacific Northwest (Ruff et al. 2026).

Because pileated woodpeckers function as ecosystem engineers strongly associated with mature forests and cavity-dependent wildlife communities, these analyses illustrate how acoustic monitoring networks developed for focal species can simultaneously generate ecologically meaningful inference for broader forest biodiversity and ecosystem structure.

Emerging analyses are also expanding the temporal and sensory dimensions of ecological inference possible from PAM datasets. Recent work using a NWFP-PAM dataset from the Olympic Peninsula demonstrated that automated classification workflows can characterize avian vocal phenology across broad environmental gradients, revealing signals associated with migration strategy, breeding chronology, and elevational change (Clapp et al. 2026). Similarly, a study of data collected in the Klamath study area examined realized acoustic niches and found that species distributions may be shaped not only by vegetation structure and climate, but also by the perceptual accessibility of acoustic information across landscapes (Habib et al. 2026).

Together, these developments illustrate how NWFP-PAM datasets are increasingly supporting analyses of temporal ecology, soundscape ecology, and sensory landscapes in addition to traditional occupancy estimation.

Recent advances in artificial intelligence are also expanding opportunities for ecological inference from archived recordings. Emerging methods using AI-assisted individual identification from vocalizations suggest that future PAM frameworks may support noninvasive approaches for studying movement, survival, territory occupancy, and demographic processes at broad spatial scales (Knight et al. 2024). Although these approaches remain under active development, they highlight the growing long-term scientific value of archived acoustic datasets and the importance of maintaining consistent, broad-scale biodiversity monitoring networks through time.

At the same time, several newly added PNW-Cnet v5 sound classes produced few or no apparent detections, likely reflecting limited training data and lower model recall rather than true biological absence. Continued development of training datasets, validation workflows, and iterative model refinement remain important priorities for the NWFP-PAM program to improve the utility of broad-scale acoustic biodiversity monitoring.

### Challenges encountered

The 2025 field season represented a substantial reduction in implementation capacity for the NWFP-PAM program relative to 2023 and 2024. Staffing limitations affecting participating federal agencies reduced the number of personnel available for field deployment, retrieval, logistics, and data processing. As a result, approximately half the number of hexagons and sampling stations were surveyed compared to the previous two years of full implementation of the NWFP-PAM network.

Although reductions in sampling effort were broadly distributed across the NWFP area, the magnitude of reductions varied among study areas depending on local staffing capacity, logistical constraints, wildfire-related access restrictions, and operational priorities. Some areas maintained near-complete sampling coverage, while others experienced substantial reductions in effort. Delayed retrievals associated with limited staffing and wildfire closures also resulted in several recording units remaining in the field beyond the intended retrieval window.

Long-term monitoring programs derive much of their scientific value from consistency in spatial coverage and repeated sampling through time. Reductions in implementation capacity can decrease precision of occupancy estimates, reduce statistical power to detect ecological change, and limit inference for smaller study areas or rare species like the northern spotted owl.

Nevertheless, the NWFP-PAM framework continued to provide broad-scale, statistically representative ecological information across much of the NWFP area during 2025 despite substantial operational constraints.

The 2025 field season also demonstrated the operational resilience of PAM approaches relative to many traditional wildlife survey methods. The ability to deploy autonomous recording units for extended sampling periods, combined with high-throughput machine learning workflows, allowed the NWFP-PAM program to maintain broad regional coverage despite substantial reductions in field staffing. Continued long-term investment in coordinated monitoring infrastructure, personnel, and analytical capacity will remain important for maintaining continuity of biodiversity monitoring under the NWFP.

### Outlook

The NWFP-PAM program continues to evolve from a northern spotted owl monitoring framework into a broader long-term biodiversity monitoring network capable of supporting multiple management and research objectives across federally administered forests in the Pacific Northwest. Continued advances in automated species identification, machine learning, and large-scale analytical workflows are increasing the efficiency and scope of ecological inference possible from PAM datasets. Increasingly, monitoring systems are functioning as integrated ecological observatories capable of supporting long-term biodiversity assessment, soundscape monitoring, phenological analysis, community ecology, and emerging forms of AI-assisted ecological inference.

Future efforts within the NWFP-PAM program will focus on improving occupancy and trend estimation for focal species, expanding biodiversity applications of the dataset, and integrating PAM-derived information with remote sensing and habitat modeling for annual distribution maps. Continued development of PNW-Cnet and associated validation workflows will further improve the utility of acoustic datasets for monitoring rare species, environmental conditions, and ecological change at regional scales. Emerging analytical approaches may also support future applications involving individual acoustic identification, behavioral inference, and noninvasive demographic monitoring using archived acoustic recordings.

Recent advances in edge computing and satellite-linked acoustic monitoring systems further illustrate the potential for future biodiversity monitoring networks capable of near real-time ecological inference and adaptive management support (Lesmeister et al. 2026a).

Autonomous monitoring systems integrating onboard convolutional neural networks, real-time species classification, and satellite transmission may improve the ability to rapidly detect invasive species, evaluate disturbance events, and support time-sensitive conservation decisions in remote landscapes. As these technologies continue to mature, future monitoring frameworks may increasingly combine long-term archival recording, real-time sensing networks, and AI-enabled analytical pipelines to support integrated ecological monitoring across large spatial scales.

Long-term continuity remains one of the most important components of the NWFP-PAM framework. As the time series expands, the scientific and management value of the monitoring network for detecting ecological change, evaluating management actions, and informing conservation planning will continue to increase. Archived acoustic recordings are likely to become increasingly valuable through time as new analytical methods emerge and additional ecological questions are developed. Maintaining broad-scale, statistically representative monitoring across the NWFP area will remain important for understanding the future trajectories of northern spotted owls, barred owls, marbled murrelets, and broader forest biodiversity under changing environmental conditions.

## Acknowledgments

Funding and support for this program was provided by: USDI Bureau of Land Management and National Park Service; USDA Forest Service Pacific Northwest and Southwest Regions, and Pacific Northwest Research Station. We are deeply indebted to the dedicated field and lab technicians that collected and processed millions of hours of bioacoustics: H. Basile, A. Habib, K. Parker, and all other crew members, collaborators, and volunteers who assisted on the project.

